# Recentrifuge: robust comparative analysis and contamination removal for metagenomics

**DOI:** 10.1101/190934

**Authors:** Jose Manuel Martí

**Affiliations:** Institute for Integrative Systems Biology (I^2^SysBio), Valencia, Spain.

**Keywords:** metagenomics, comparative analysis, contamination, low biomass, robustness, Centrifuge, LMAT, CLARK

## Abstract

Metagenomic sequencing is becoming widespread in biomedical and environmental research, and the pace is increasing even more thanks to nanopore sequencing. With a rising number of samples and data per sample, the challenge of efficiently comparing results within a specimen and between specimens arises. Reagents, laboratory, and host related contaminants complicate such analysis. Contamination is particularly critical in low microbial biomass body sites and environments, where it can comprise most of a sample if not all. Recentrifuge implements a robust method for the removal of negative-control and crossover taxa from the rest of samples. With Recentrifuge, researchers can analyze results from taxonomic classifiers using interactive charts with emphasis on the confidence level of the classifications. In addition to contamination-subtracted samples, Recentrifuge provides shared and exclusive taxa per sample, thus enabling robust contamination removal and comparative analysis in clinical and environmental metagenomics.

**Author summary:** Whether in a clinical or environmental sample, metagenomics can reveal what microorganisms exist and what they do. It is indeed a powerful tool for the study of microbial communities which requires equally powerful methods of analysis. Current challenges in the analysis of metagenomic data include the comparative study of samples, the degree of uncertainty in the results, and the removal of contamination. The scarcer the microbes are in an environment, the more essential it is to have solutions to these issues. Examples of sites with few microbes are not only habitats with low levels of nutrients, but also many body tissues and fluids. Recentrifuge’s novel approach combines statistical, mathematical and computational methods to tackle those challenges with efficiency and robustness: it seamlessly removes diverse contamination, provides a confidence level for every result, and unveils the generalities and specificities in the metagenomic samples.

## Introduction

Studies of microbial communities by metagenomics are becoming more and more popular in different biological arenas, like environmental, clinical, food and forensic studies (Miller et al. 2013; Ercolini 2013; Fricke et al. 2011). New DNA and RNA sequencing technologies are boosting these works by dramatically decreasing the cost per sequenced base. Scientists can now analyze sets of sequences belonging to microbial communities from different sources and times to unravel longitudinal —spatial or temporal— patterns in the microbiota (see Supplemental Figure 5 for an example model). In shotgun metagenomic sequencing (SMS) studies, researchers extract and purify nucleic acids from each sample, sequence them, and analyze the sequences through a bioinformatics pipeline (see Supplemental Figure. 6 and 7 for detailed examples). With the development of nanopore sequencing, portable and affordable real-time SMS is a reality (Edwards et al. 2016).

### Contamination in metagenomics

In the case of low microbial biomass samples, there is very little native DNA from microbes; the library preparation and sequencing methods will return sequences whose principal source is contamination (Weiss et al. 2014; Kim, Hofstaedter, et al. 2017). Sequencing of RNA requiring additional steps introduces still further biases and artifacts (Kim, Song, et al. 2016), which in case of low microbial biomass studies translates into a severe problem of contamination and spurious taxa detection (Perlejewski et al. 2016). The clinical metagenomics community is stressing the importance of negative controls in metagenomics workflows and, recently, raised a fundamental concern about how to subtract the contaminants from the results (Ruppé and Schrenzel 2018).

From the data science perspective, this is just another instance of the importance of keeping a good *signal-to-noise ratio* (Skolnik 2008). When the *signal* (inherent DNA/RNA, target of the sampling) approaches the order of magnitude of the *noise* (acquired DNA/RNA from contamination and artifacts), particular methods are required to tell them apart.

The roots of contaminating sequences are diverse, as they can be traced back to nucleic acid extraction kits —the *kitome*— (Salter et al. 2014; Thoendel et al. 2017), reagents and diluents (Olm et al. 2017; Kulakov et al. 2002), the host (Ames, Gardner, et al. 2015) and the post-sampling environment (Lusk 2014), where contamination arises from different origins such as airborne particles, crossovers between current samples or DNA remains from past sequencing runs (Gruber 2015). Variable amounts of DNA from these sources are sequenced simultaneously with native microbial DNA, which could lead to severe bias in magnitudes like abundance and coverage, particularly in low microbial biomass situations (Nayfach and Pollard 2016). If multiplex sequencing uses simple-indexing, false assignments could be easily beyond acceptable rates (Kircher et al. 2012). Even the metagenomic reference databases have a non-negligible amount of cross-contamination (Ames, Gardner, et al. 2015; Gruber 2015; Lu and S Salzberg 2018).

Regarding the *kitome*, it varies even within different lots of the same products. For example, the DNeasy PowerSoil Kit (formerly known as PowerSoil DNA Isolation Kit), a product that usually provides significant amounts of DNA and has been widely used, including Earth Microbiome Project and Human Microbiome Project, often yields a background contamination by no means negligible (Kim, Hofstaedter, et al. 2017). The lower the biomass in the samples, the more essential it is to collect negative control samples to help in the contamination background assessment because, without them, it would be almost impossible to distinguish inherent microbiota in a specimen —signal— from contamination —noise—.

Assuming that the native and contaminating DNA are accurately separated, the problem of performing a reliable comparison between samples remains. In general, the taxonomic classification engine assigns the reads from a sequencing run to different taxonomic ranks, especially if the method uses a more conservative approach like the lowest common ancestor —LCA— (Ames, Hysom, et al. 2013). While LCA drastically reduces the risk of false positives, it usually spreads the taxonomic level of the classifications from the more specific to the more general. Even if the taxonomic classifier does not use the LCA strategy, each read is assigned a particular score or confidence level, which should be taken into account by any downstream application as a reliability estimator of the classification.

On top of these difficulties, it is still more challenging to compare samples with very different DNA yields, for instance, low microbial biomass samples versus high biomass ones, because of the different resolution in the taxonomic levels. This sort of problem also arises when the samples, even with DNA yields in the same order of magnitude, have an entirely different microbial structure so that the minority and majority microbes are fundamentally different between them (Nayfach and Pollard 2016). Finally, a closely related problem emerges in metagenomic bioforensic studies and environmental surveillance, where it is essential to have a method prepared to detect the slightest presence of a particular taxon (Bazinet et al. 2017; Doggett et al. 2016; Fricke et al. 2011) and provide quantitative results with both precision and accuracy.

### Comparison and validation of metagenomic results

From the beginning, the application of SMS to environmental samples supplied biologists with an insight of microbial communities not obtainable from the sequencing of Bacterial Artificial Chromosome (BAC) clones or 16S rRNA (Allen et al. 2004; Venter et al. 2004). The scientific community soon underlined the need and challenges of comparative metagenomics (Schloss and Handelsman 2005; Tringe et al. 2005). MEGAN (Huson, Auch, et al. 2007), one of the first metagenomic data analysis tools, provided in its initial release a very basic comparative of samples, which has improved with an interactive approach in more recent versions (Huson, Beier, et al. 2016). In general, metagenomic classification and assembly software is more intra-than inter-sample oriented (Breitwieser, Lu, et al. 2017). Several tools have tried to fill the gap, starting with CoMet (Lingner et al. 2011), a web-based tool for comparative functional profiling of metagenomes. Soon later, a different approach appeared with the discovery of the crAssphage thanks to the crAss software (Dutilh et al. 2012), which provides reference-independent comparative metagenomics using cross-assembly. The following year, a new tool for visually comparing microbial community structure across microbiomes, Community-analyzer (Kuntal et al. 2013), arrived. In 2014, yet another alternative came, COMMET (Maillet et al. 2014), a piece of software that goes a step further by enabling the combination of multiple metagenomic datasets. Two years later, a package for direct comparative metagenomics by using k-mers (Benoit et al. 2016) was published.

In 2015, a highly publicized report on the metagenomics of the New York subway suggested that the plague and anthrax pathogens were part of the normal subway microbiome. Soon afterward, several critics arose (Ackelsberg et al. 2015) and, later, reanalysis of the New York subway data with appropriate methods did not detect the pathogens (Tiffany et al. 2016). As a consequence of this and other similar problems involving metagenomic studies, a work directed by Rob Knight (A González et al. 2016) emphasized the importance of validation in metagenomic results and issued a tool based on BLAST (Platypus Conquistador). This software confirms the presence or absence of a taxon of interest within SMS datasets by relying on two reference sequence databases: one for inclusions, with the sequences of interest, and the other for exclusions, with any known sequence background. Another BLAST-based method for validating the assignments made by less precise sequence classification programs has been recently announced (Bazinet et al. 2017).

Recentrifuge approach to increased confidence in the results of taxonomic classification engines follows a dual strategy. Firstly, it accounts for the score level of the classifications in every single step. Secondly, it uses a robust contamination removal algorithm that detects and selectively eliminates various types of contaminants, including crossovers. Recentrifuge currently supports two high-performance taxonomic classifiers, Centrifuge (Kim, Song, et al. 2016) and LMAT (Ames, Hysom, et al. 2013). The interactive interface of Recentrifuge enables researchers to analyze results from those classifiers using scored Krona-like charts (Ondov et al. 2011). In addition to the plots for the raw samples, Recentrifuge generates four different sets of scored charts for each taxonomic level of interest: control-subtracted samples, shared taxa (with and without subtracting the controls), and exclusive taxa per sample. This battery of analysis and plots permits robust comparative analysis of multiple samples in metagenomic studies, especially useful in case of low microbial biomass environments or body sites.

## Methods

### Visualization and computing kernel

The visualization part of Recentrifuge is based on interactive hierarchical pie charts. The Krona (Ondov et al. 2011) 2.0 JavaScript library developed at the Battelle National Biodefense Institute was adapted and extended to fit the particular needs of the problem.

The computing kernel of Recentrifuge is a metagenomic data analysis engine written in Python. Recentrifuge’s novel approach combines robust statistics, arithmetic of scored taxonomic trees, and parallel computational algorithms to achieve its goals.

For each sample, according to the NCBI Taxonomy, Recentrifuge populates a logical taxonomic tree, with the leaves usually belonging to the lower taxonomic levels like varietas, subspecies or species. The methods involving trees were implemented as recursive functions for compactness and robustness, making the code less error-prone. One of such methods is essential for understanding the way Recentrifuge prepares samples before any comparison or operation, like control subtraction. It recursively “folds the tree” for any sample if the number of assigned reads to a taxon is under the mintaxa setting (minimum reads assigned to a taxon to exist in its own), or because the taxonomic level of interest is over the assigned taxid (taxonomic identifier). See Figure 1A for a working example of the method in action for two samples. The same procedure applies to the trees of every sample in the dataset. This method does not just “prune the tree”, on the contrary, it accumulates the counts *n*_*i*_ of a taxon in the parent ones *n*_*p*_ and recalculates the parent score *σ*_*p*_ as a weighted average taking into account the counts and score of both. In general, the new score of parent taxa, 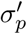 is calculated as follows:

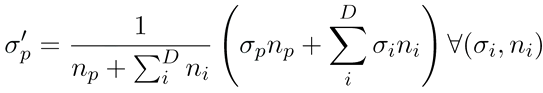

where 0 < *ni* < mintaxa and D is the number of descendant taxa that are to be accumulated in the parent one and σ_*i*_ their respective scores. This is done recursively until the desired conditions are met. This method is applied, at a given taxonomic level, to the trees of every sample before being compared in search for the shared and exclusive taxa. The mintaxa parameter defaults to 10 as in LMAT, but this value can be modified and set independently for control and real samples.

**Figure 1.**
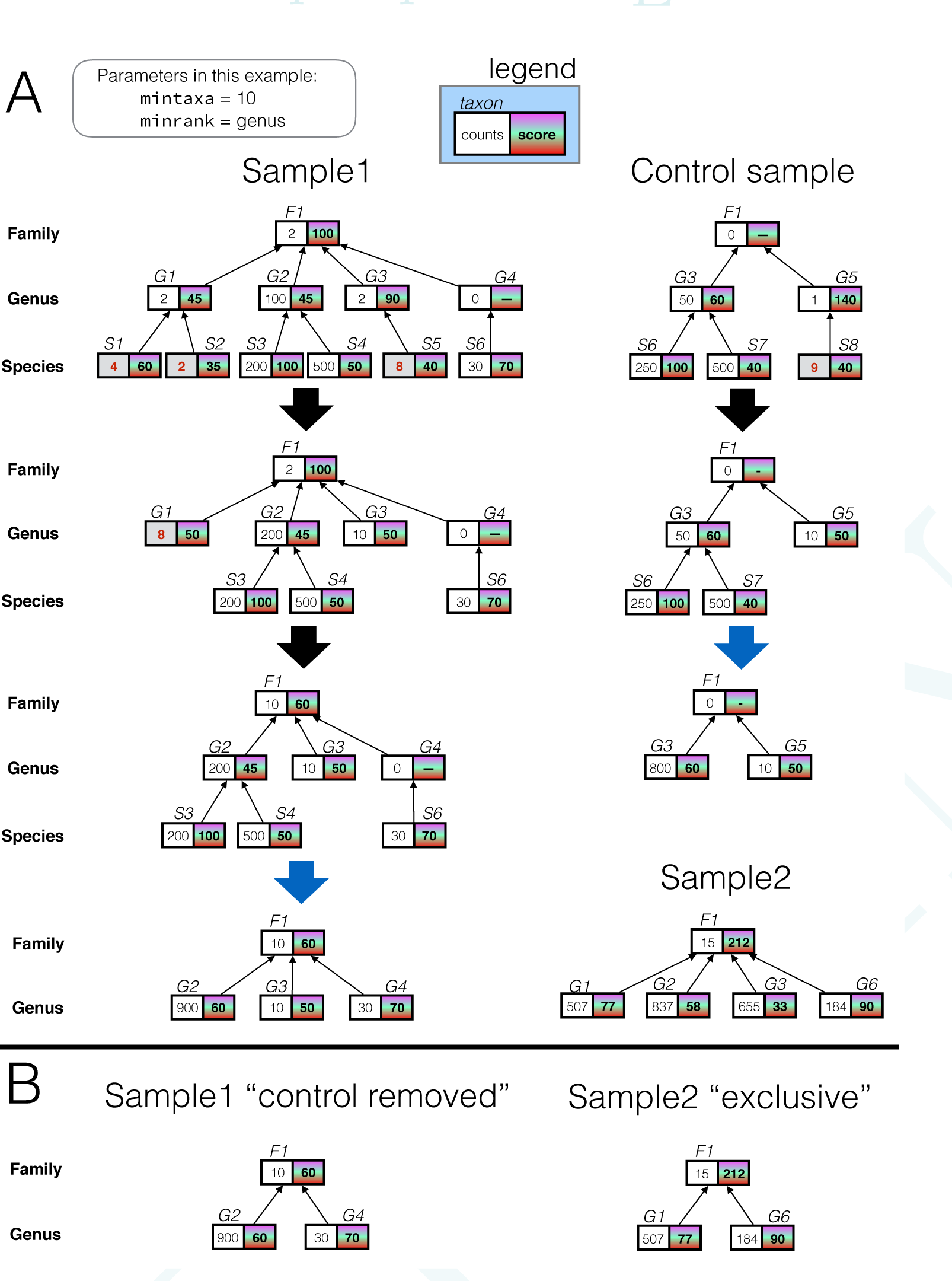
Operating with taxonomic trees. (**A**) Example of the recursive function which “folds the tree” to prepare the taxonomic trees for further operations, with the parameter mintaxa set to 10 (default) and the minimum rank of interest minrank set to genus. Initially, their trees show the direct taxonomic classification results. Then, recursively, the leaves of the tree are accumulated in the parent node if their number of assigned reads is under mintaxa (shown in red and bold counts) or if their corresponding taxonomic rank is below minrank. In this ‘folding’ the parent score is updated with a weighted average of its own score and the ones of the descendants that are being accumulated. E.g., after the 1st step, the G1 taxon at the sample is updated with *n*_*p*_*=2+4+2=8* counts and score of 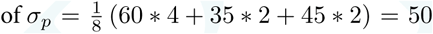. As the counts for G1 are still under mintaxa, in the 2nd step they are accumulated in F1 and its score updated to 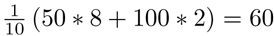. (**B**) Continuing with the example in (A), at genus level, there are two derived samples: the right one with the control removed from *Samplei*, the left one with the exclusive taxa of *Sample2* (those taxa not present in the rest of samples, in this case, the control and *Sample1)*.

For implementation details of the Recentrifuge computing kernel please see Supplementary Section 2.

### Derived samples

In addition to the input samples, Recentrifuge includes some sets of derived samples in its output. After parallel calculations for each taxonomic level of interest, it adds hierarchical pie plots for CTRL (control subtracted), but also for EXCLUSIVE, SHARED and SHARED_CONTROL samples, defined below.

Let 𝕋 mean the set of taxids in the NCBI Taxonomy and *T*_*s*_ the collection of taxids present in a sample s.

If *R*_*s*_ stands for the set of reads of a sample *s* and *C*_*s*_ for the group of them classifiable, then the taxonomic classification c is a function from *C*_*s*_ to 𝕋, i.e., *C*_*s*_ to 𝕋, i.e., 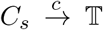, where 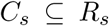 and *c*[*C*_*s*_] = 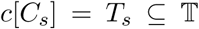 The set *L* of the 30 different taxonomic levels used in the NCBI is ordered in accordance with the taxonomy, so (*L*, <) is a strictly ordered set, since *forma < varietas < subspecies < · · · < domain*. Then, 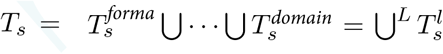, where 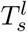 represents the collection of taxa belonging to a sample s for a particular taxonomic rank or level l. Related with this, we can write as 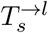 the taxa of the sample s for a taxonomic level l once we have applied the ‘tree folding’ to such level l detailed in the previous subsection —and Figure iA.

For a taxonomic rank *k* of interest, in a series of S samples where there are *N < S* negative controls, Re -centrifuge computes the sets of taxa in the derived samples CTRL (^CTRL^*T^K^_S_*), EXCLUSIVE (^EXCL^*T^K^_S_*), SHARED (^SHARED^*T^K^*) and SHARED_CONTROL (^SHARED^-^CTRL^*T^K^*) as:

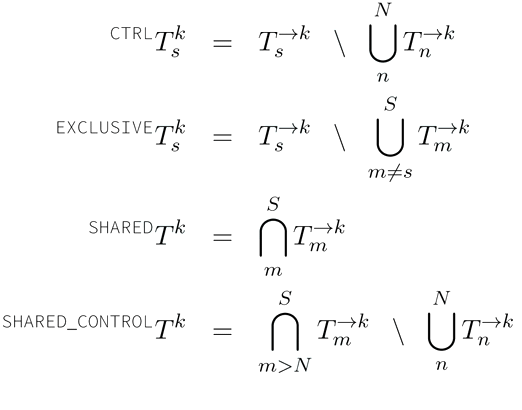

Please see Figure 1B for examples. Finally, Recentrifuge generates in parallel a set of SUMMARY samples condensing the results for all the taxonomic levels of interest.

### Robust contamination removal

For a taxonomic rank *k*, after the ‘tree folding’ procedure detailed above, the contamination removal algorithm retrieves the set of candidates 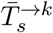 to contaminant taxa from the *N* < *S* control samples. Depending on the relative frequency (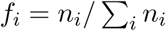) of these taxa in the control samples and if they are also present in other specimens, the algorithm classifies them in contamination level groups: critical, severe, mild and other. Except for the latter group, the contaminants are removed from non-control samples. Then, Recentrifuge checks any taxon in the other contaminants group for crossover contamination so that it eliminates any taxon marked as a crossover from every sample except the one or ones selected as the source of the pollution. In detail, the algorithm removes any taxon 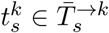 from a non-control sample unless it passes the robust crossover check: a statistical test screening for overall outliers and an order of magnitude test against the control samples. See Figure 2 for an example of this procedure.

**Figure 2.**
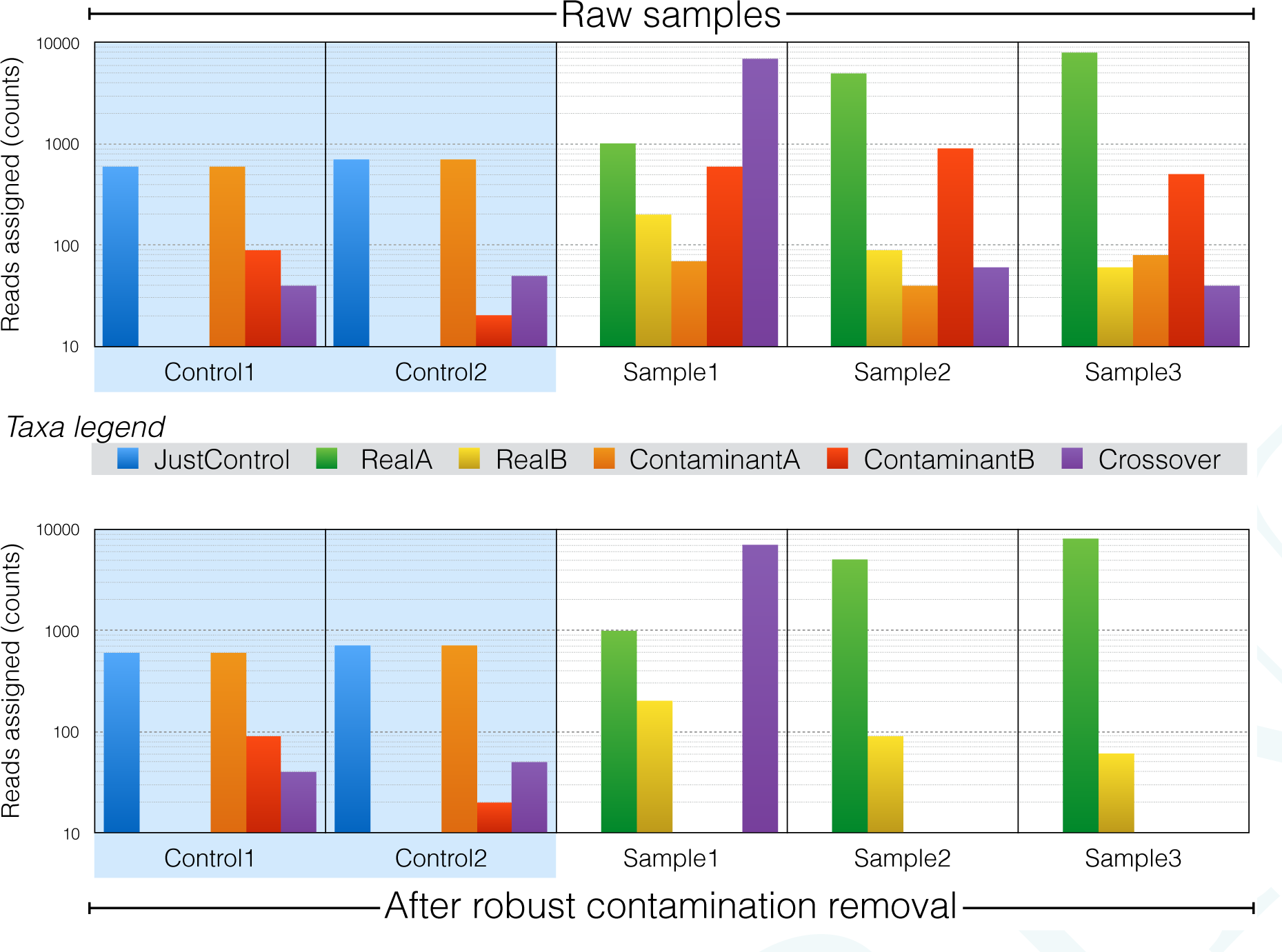
Robust contamination removal. This is a hypothetic example with 5 samples and 6 dominant taxa to illustrate how the algorithm works. The top and bottom part of the figure shows the absolute frequency of reads assigned to the taxa before and after the contamination removal, respectively. There are two control samples, not modified throughout the process. In the rest of specimens, the general contaminants (those taxa present in the controls and other samples, like *ContaminantA* and *ContaminantB)* are removed, except in case of crossover contamination: those taxa are kept in the source(s) sample(s) —Sample1 here— and removed from other real samples (Sample2 and Sample3 in this example).

The robust crossover tests are defined as follows:

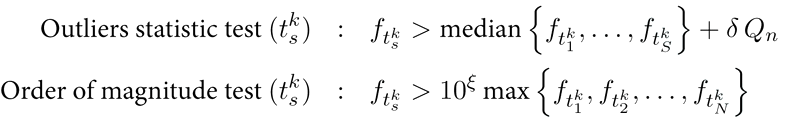

where *Q*_*n*_ (Rousseeuw and Croux 1993) is a scale estimator to be discussed below, and *δ* and *ξ* are constant parameters of the robust contamination removal algorithm. The parameter *δ* (3 to 5 by default) is an outliers cutoff factor, while *ξ* (2 to 3 by default) is setting the difference in order of magnitude between the relative frequency of the candidate to crossover contaminator in the sample s and the greatest of such values among the control samples.

*Q*_*n*_ is the chosen scale estimator for screening the data for outliers because of his remarkably general robustness and other advantages compared to other estimators (Rousseeuw and Croux 1993; Rousseeuw and Croux 1994), like the MAD (median absolute deviation) or the *k*-step *M*-estimators. It has a 50% breakpoint point, a smooth influence function, very high asymptotic efficiency at Gaussian distributions and is suitable for asymmetric distributions, which is our case, all at a reasonable computational complexity, as low as *O(n)* for space and *O(n* log *n*) for time. So, here:

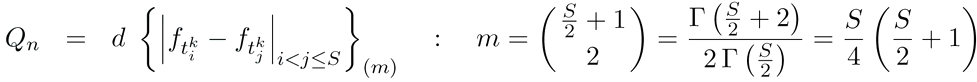

where *d* = 3.4760 is a constant selected for asymmetric non-gaussian models similar to the negative exponential distribution, *m* refers to the *m*th order statistics of the pairwise distances and Γ is the Gamma function.

## Results

Recentrifuge is a metagenomics analysis software with two different main parts: the computing kernel, implemented and parallelized from scratch using Python, and the interactive interface, written in JavaScript as an extension of the Krona (Ondov et al. 2011) JS library to take full advantage of the classification confidence level. Figure 3 summarizes the package context and data flows. Recentrifuge straightforwardly accepts Centrifuge (Kim, Song, et al. 2016), LMAT (Ames, Hysom, et al. 2013), and CLARK(S) (Ounit and Lonardi 2016) output files, thus enabling scored oriented analysis and visualization for these tools. Recentrifuge also supports LMAT plasmids assignment system (Ames, Gardner, et al. 2015).

**Figure 3.**
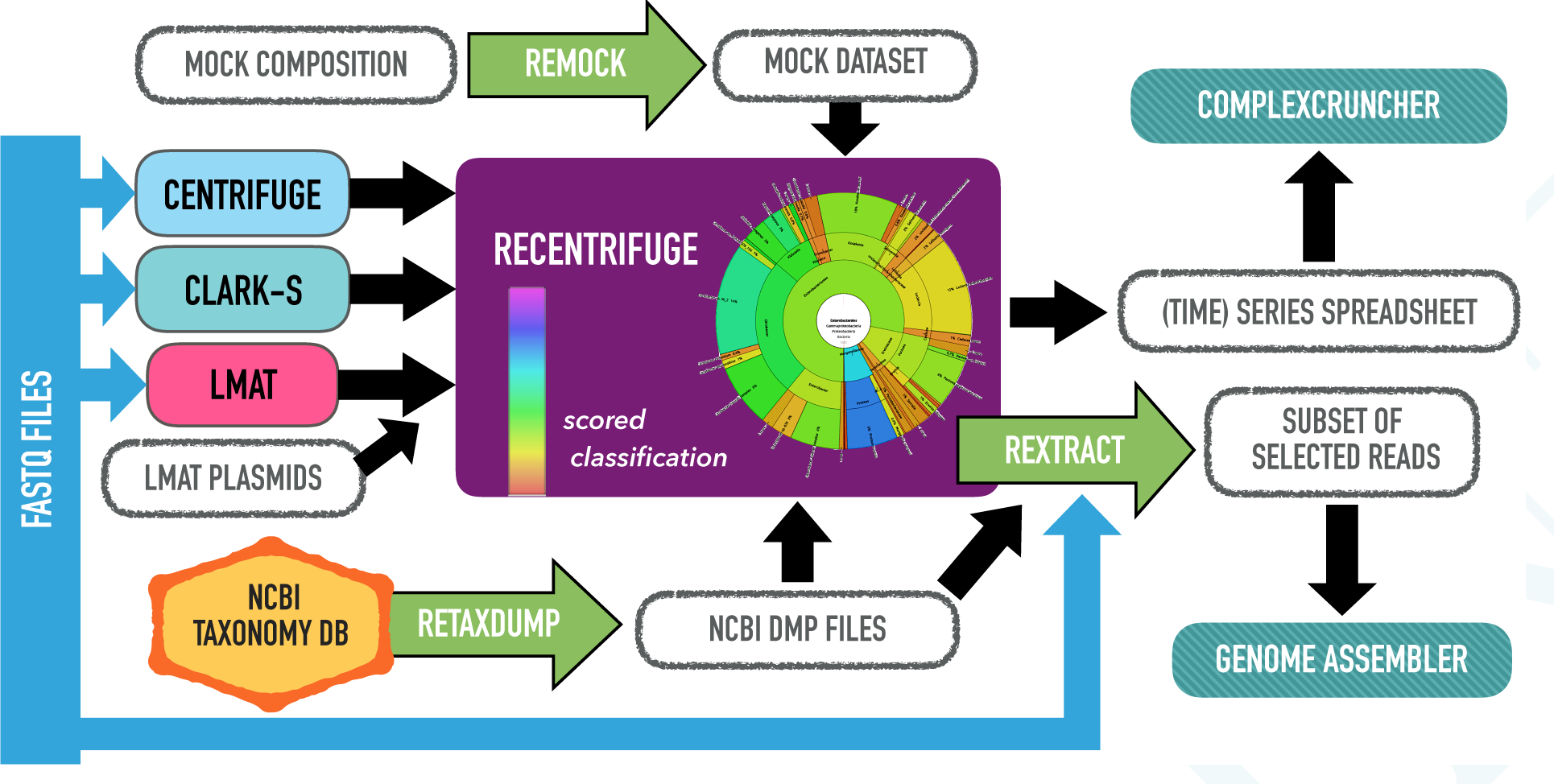
Outline of the Recentrifuge package with its context and main data flows. The green arrows indicate pipelines implemented by means of additional scripts. Recentrifuge accepts both Centrifuge (Kim, Song, et al. 2016), LMAT (Ames, Hysom, et al. 2013), and CLARK-S (Ounit and Lonardi 2016) direct output files, thus enabling a scored oriented visualization for any of these codes. Recentrifuge is also supporting LMAT plasmids assignment system (Ames, Gardner, et al. 2015). The additional output of Recentrifuge to different text field formats enable further longitudinal (time or space) series analysis, for example, using *cmplxCruncher* (in development). The NCBI Taxonomy dump databases are easily retrieved using *Retaxdump* script. *Rextract* utility extracts a subset of reads of interest from single or paired-ends FASTQ input files, which can be used in any downstream application, like genome assembling and visualization. *Remock* easily creates mock Centrifuge samples, useful for validation and for including previously known contaminants.

To ensure the broadest portability for the interactive visualization of the results, the central outcome of Recentrifuge is a stand-alone HTML file which can be loaded by any JavaScript-enabled browser. Figure 4 shows a labeled screenshot of the corresponding Recentrifuge web interface for an example of SMS study (see Supplemental Figure 5). The package also provides statistics about the reads and their classification performance. Another Recentrifuge output is an Excel spreadsheet detailing all the classification results in a compact way. This format is adequate for further data mining on the data, for example, as input for applications such as longitudinal (time or space) series analyzers like *cmplxCruncher* (in development). Besides, the user can choose between several scoring schemes algorithms, some of which are more interesting for datasets containing variable length reads, for example, the ones generated by nanopore sequencers. Finally, different filters are available, like the ones for selective exclusion or inclusion of entire clades, or the minimum score threshold. This latter filter can be used to generate different output sets from a single run of the classifier with a low minimum hit length (MHL) setting. This strategy not only saves time and computational resources but also makes compatible the advantage of a low MHL in low quality reads, on the one hand, with the benefit of a high MHL to avoid false positives, on the other hand.

**Figure 4.**
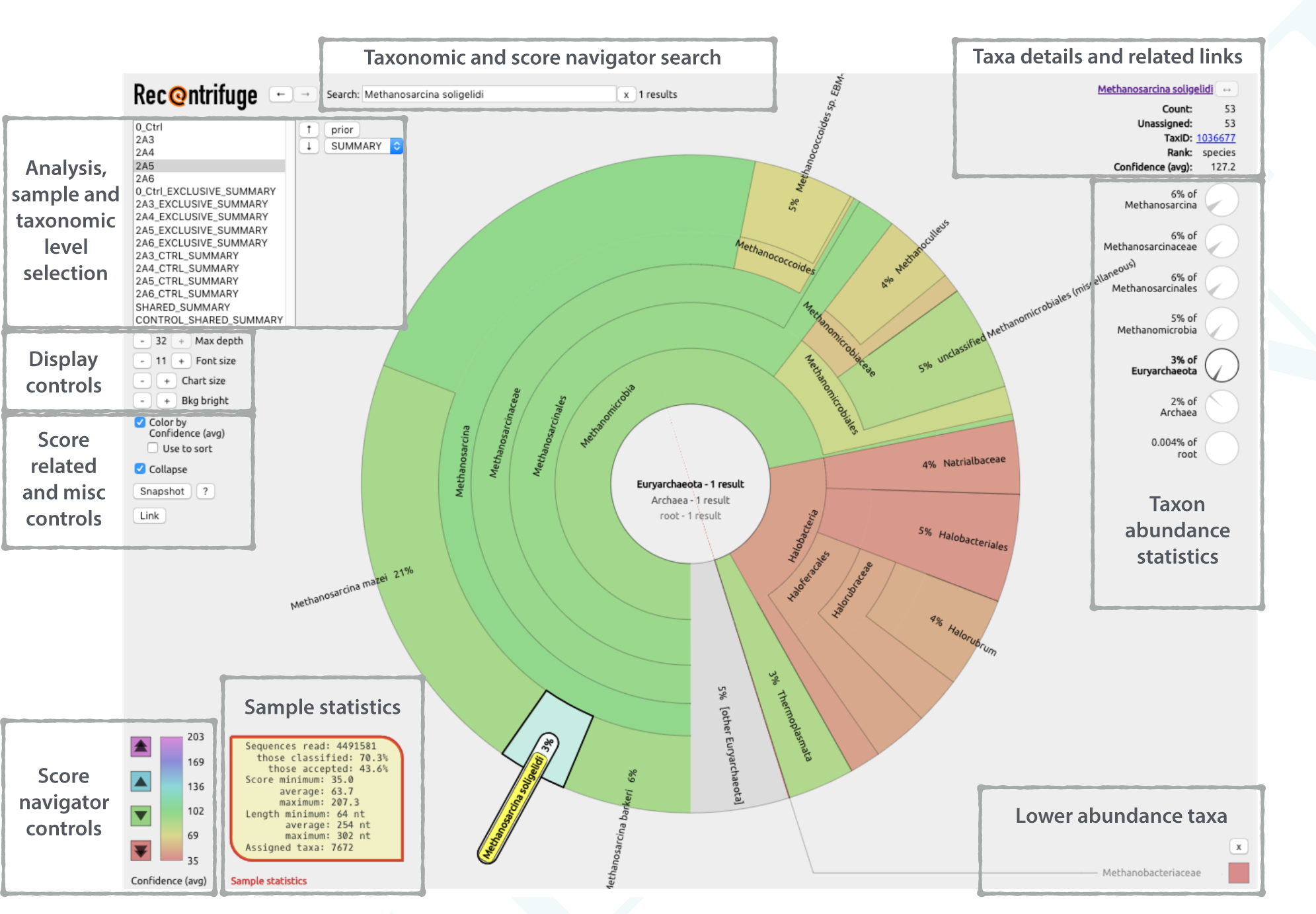
Layout of the Recentrifuge interface. This figure is an explained screenshot of the Recentrifuge web interface for an SMS study (see Supplemental Figure 5 for details of the example). It highlights the principal parts of the interface, which are also labeled. The sample 2A5 was selected (see the sample selection box in the top left under the Recentrifuge logo), so the key statistics for this sample appeared in the bottom left of the view. In the center, there was the corresponding hierarchical pie chart, with zoom in the phylum Euryarchaeota. For each taxon, the background color reflected the average confidence level of the taxonomic classification following the scale plotted in the bottom left of the figure, where there were also buttons for the score navigator. Since the interface had the option disabled, Recentrifuge did not sort the taxa attending to the average confidence level. In this particular case, the taxon *Methanosarcina soligelidi* was selected in the pie chart, thus prompting the display of taxon-related statistics and links in the top right of the figure. The current links were to Google Scholar and NCBI Taxonomic Browser. The statistics included: the number of reads assigned to this or lower taxonomic levels (Count) and their average confidence (Confidence —avg—), the number of reads just assigned to this level (Unassigned), the NCBI taxid (TaxID) and rank (Rank), and some information about relative frequencies.

Further products are obtained using auxiliary tools in the package, like *Rextract*, a script which helps in extracting a subset of classified reads of interest from the single or paired-ends FASTQ input files. These reads can be used in any downstream application, such as genome visualization and assembling. *Remock* is a script for easily creating mock Centrifuge samples, which is useful not only for testing and validation purposes, but also for introducing a list of previously known contaminants to be taken into account by the robust contamination removal algorithm.

## Discussion

Recentrifuge enables robust contamination removal and score-oriented comparative analysis of multiple samples, especially in low microbial biomass metagenomic studies, where contamination removal is a must. In any SMS study with related samples, including negative controls, Recentrifuge generates four additional sets of scored charts: the samples with the contamination subtracted, the exclusive taxa per sample, and the shared taxa with and without control taxa subtracted (see Supplemental Figure 8).

Just as it is important to accompany any physical measurement by a statement of the associated uncertainty, it is desirable to accompany any read classification by a confidence estimation of the taxon that has been finally assigned. Recentrifuge reads the different scores given by Centrifuge, LMAT, or CLARK(S) to any classified read and uses this valuable information to calculate an averaged confidence level for each classification node. Depending on the scoring scheme selected by the user, this value maybe also a function of further parameters, such as read length. This scheme is especially useful in the case of significant variations in the length of the reads, like in the datasets generated by nanopore sequencers.

Only a few codes, such as Krona (Ondov et al. 2011) and MetaTreeMap (Hebrard and Taylor 2016), are hitherto able to handle a score assigned to the classification nodes. In Recentrifuge, the calculated score propagates to all the downstream analysis and comparisons, including the interface, an interactive framework for an easy assessment of the validity of the taxonomic assignments. That is an essential advantage of Recentrifuge over other metagenomic dataset analysis tools.

Thanks to the robust contamination removal and the score-oriented comparative analysis of multiple samples in metagenomics, Recentrifuge can play a key role not only in the study of oligotrophic microbes in environmental samples but also in a more reliable detection of minority organisms in clinical or forensic samples.

## Availability

Recentrifuge’s main website is www.recentrifuge.org. It is also available in Github (https://github.com/khyox/re-centrifuge) under different free software licenses. The wiki (https://github.com/khyox/recentrifuge/wiki) is the most extensive and updated source of documentation for Recentrifuge.

## Acknowledgments

I would like to thank Jordi Burguet-Castell and Andrea Salvador-Pascual for critical comments on the manuscript. I would also like to thank Carlos P. Garay and Jose Fco. Martí for valuable conversations about Recentrifuge.

## Supplemental Material of Recentrifuge: robust comparative analysis and contamination removal for metagenomic data

## 1 Longitudinal metagenomics

In longitudinal metagenomics, scientists retrieve and analyze sets of sequences belonging to microbial communities from different sources, times, patients, or body sites to unravel spatial, temporal or clinical patterns in the microbiota (see Supplemental Figure 5 for an example). In those studies involving shotgun metagenomic sequencing (SMS), researchers extract DNA/RNA from each sample (using a commercial kit or an optimized custom protocol), sequence the purified DNA/RNA and analyze the reads through a bioinformatics pipeline. Supplemental Figure 6 summarizes this protocol and Supplemental Figure 7 details the core of the analysis. Supplemental Figure 8 summarizes the instant benefits of introducing Recentrifuge in an SMS study.

**Figure 5.**
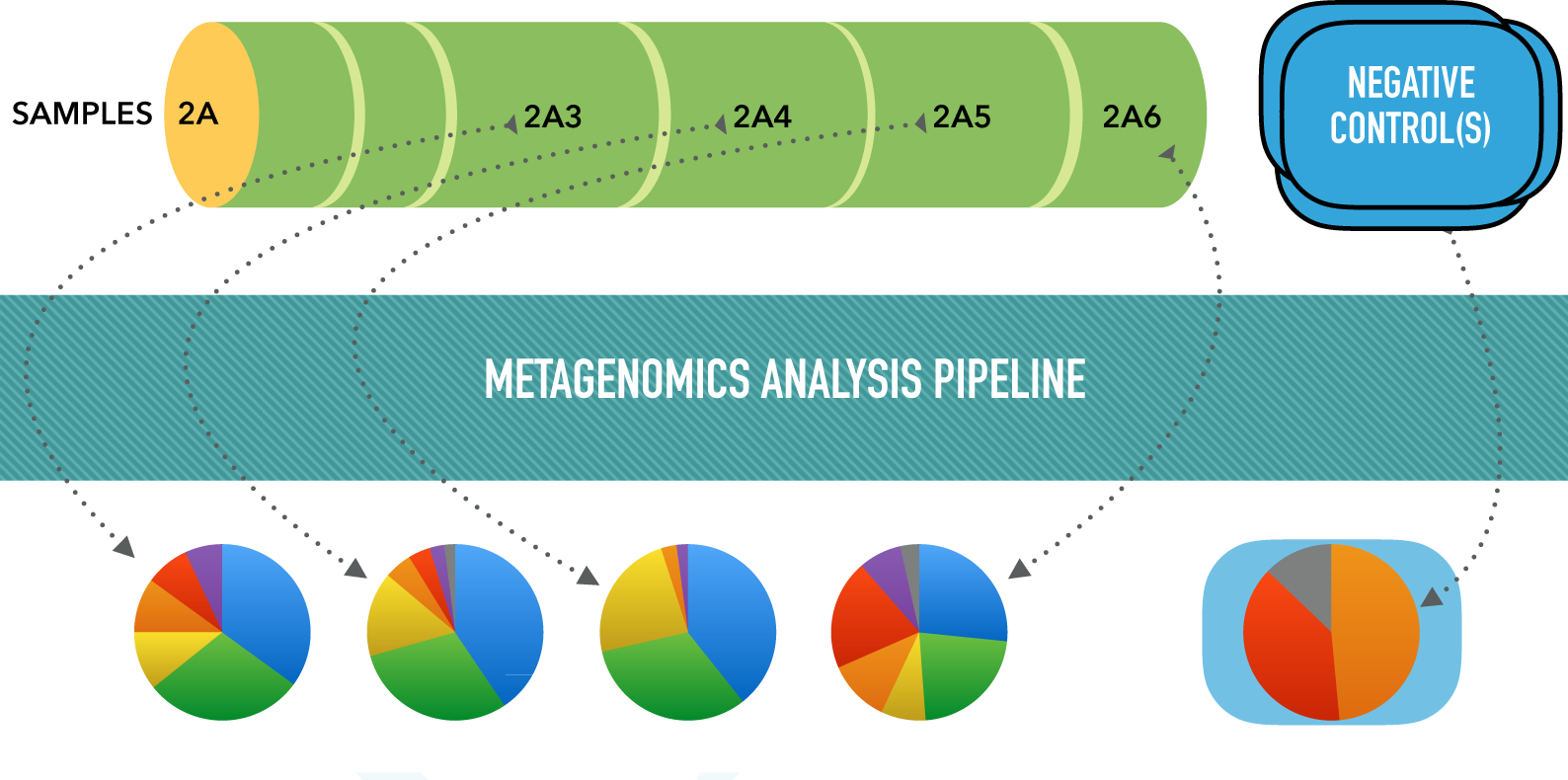
(Supplemental) Example of longitudinal SMS study. This is an example outlining the problem of comparing different but related samples in a SMS study. The sample named 2A is subdivided longitudinally into six subsamples whose DNA/RNA is extracted along with negative control samples. The purified DNA/RNA is then sequenced, and the generated sequencing reads are processed through a metagenomics analysis pipeline (as the one detailed in Supplemental Figure 6). A collection of different datasets are finally produced, which should be adequately compared to elucidate lengthwise patterns in the microbiota within the 2A sample.

**Figure 6.**
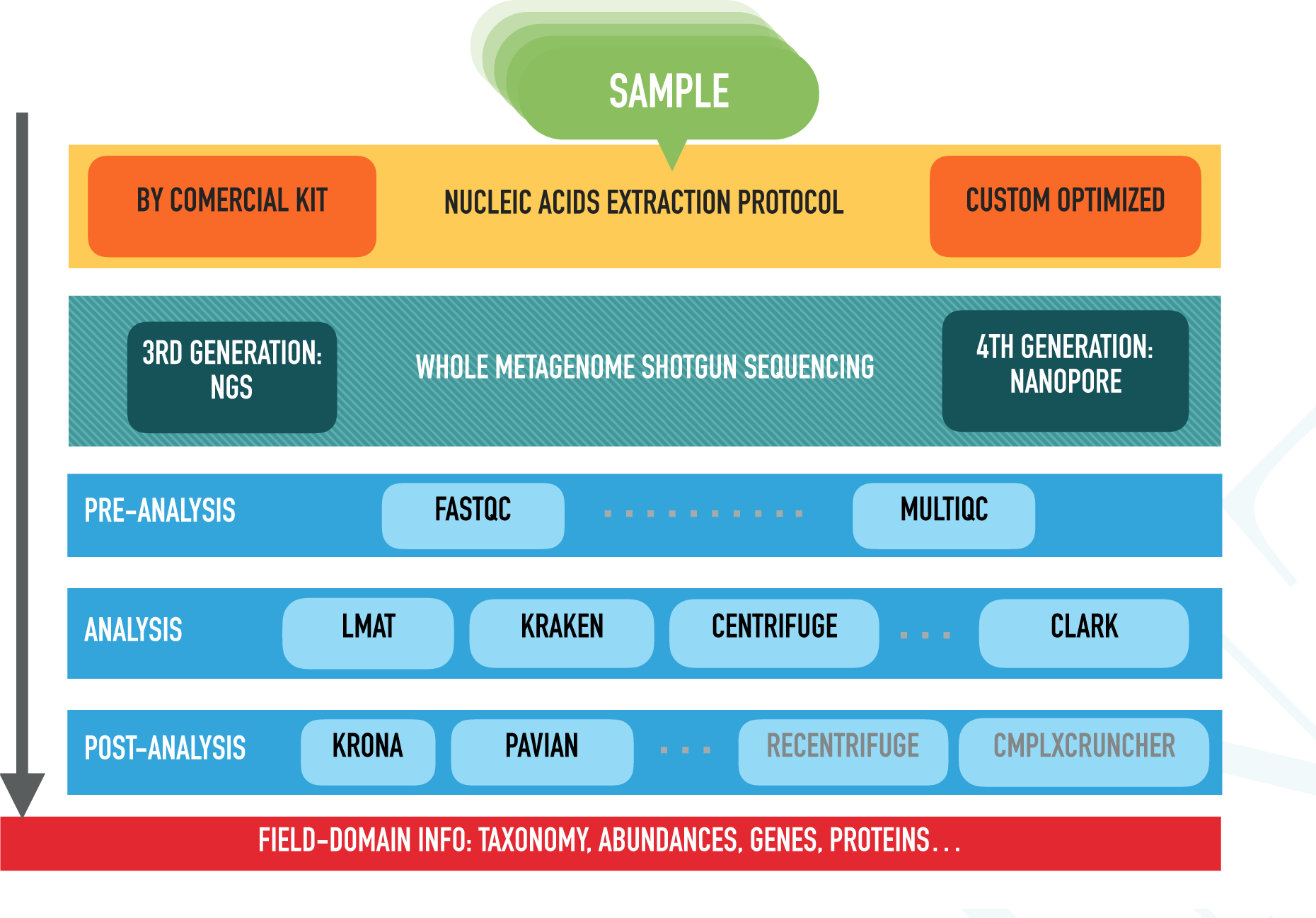
(Supplemental) Typical steps of an SMS study. A study involving SMS, like the one illustrated by Supplemental Figure 5, spans in a number of stages to extract valuable field-domain information starting from the original samples. For each specimen, the researcher extracts DNA/RNA using a commercial kit, a custom protocol optimized for the type of sample, or a combination of both. Next, a technician prepares a library matching the target sequencing technology with the purified DNA/RNA, which is then sequenced. A bioinformatics pipeline processes the reads that the sequencer provides. We could roughly separate such process in three consecutive steps. First, in the pre-analysis, codes like FastQC (Babraham Bioinformatics, 2016) and MultiQC (Ewels et al. 2016) quality-check the reads. Second, in the analysis stage, the most computationally intensive one, software packages like LMAT (Ames, Hysom, et al. 2013), Kraken (Wood and SL Salzberg 2014), CLARK (Ounit, Wanamaker, et al. 2015), Centrifuge (Kim, Song, et al. 2016), and CLARK-S (Ounit and Lonardi 2016) —see Supplemental Figure 7 for details— classify the reads taxonomically or functionally. Finally, in the post-analysis step, different tools like Krona (Ondov et al. 2011), Pavian (Breitwieser and SL Salzberg 2016), or Recentrifuge further process the results to enable more in-depth analysis and improved visualization.

**Figure 7.**
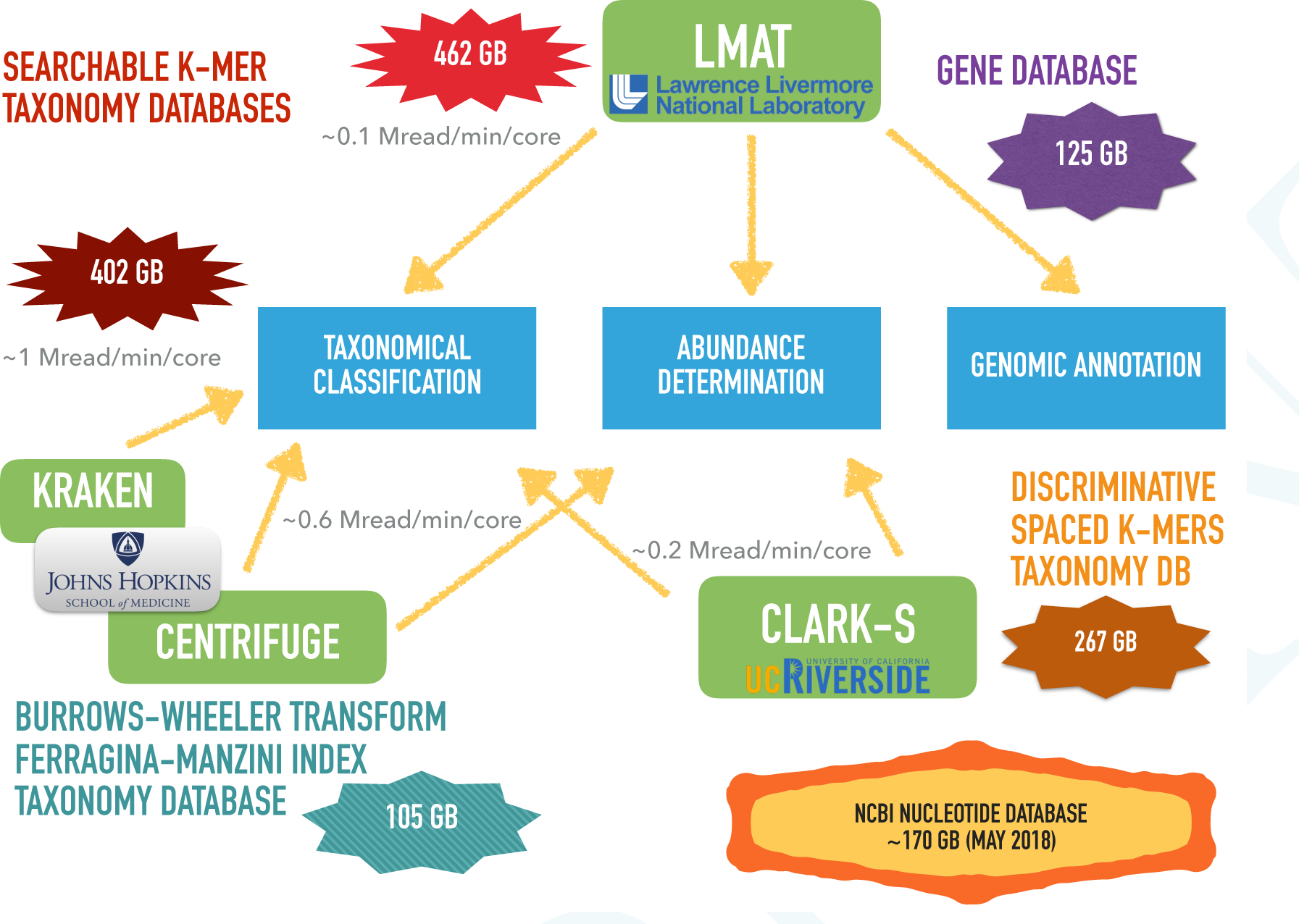
(Supplemental) Computational analysis in an SMS study. These are intensive codes in both CPU and memory (sometimes, they are input/output intensive too), like LMAT (Ames, Hysom, et al. 2013), Kraken (Wood and SL Salzberg 2014) and, more recently, CLARK-S (Ounit and Lonardi 2016) and Centrifuge (Kim, Song, et al. 2016). All these tools are performing taxonomic classification and abundance estimation, whereas LMAT is also able to annotate genes. For the taxonomic classification, both LMAT and Kraken use an exact k-mer matching algorithm with large databases (~ 100 GiB) while Centrifuge use compression algorithms to reduce the databases size (~ 10GiB) but at some speed expense. CLARK-S use discriminative spaced k-mers to improve the sensitivity but with a toll on the performance. The most complete LMAT database is approaching half terabyte of required memory while the Centrifuge database generated in-house in March 2018 from the NCBI Nucleotide (NCBI 1988a) *nt* database (~ 170 GiB) occuped just 105 GiB. The equivalent spaced k-mers database of CLARK-S generated in May 2018 taked 267 GiB of disk space.

**Figure 8.**
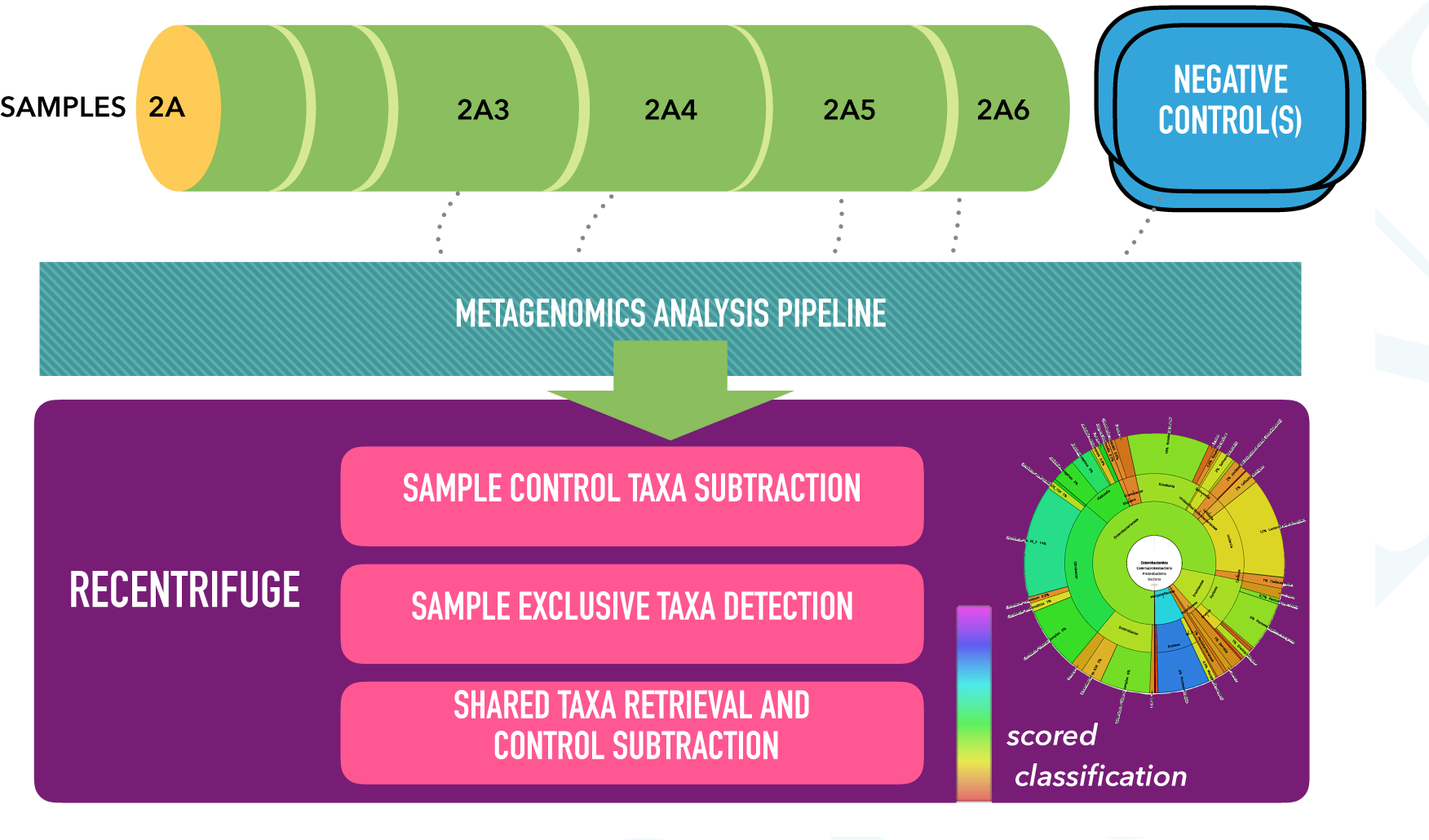
(Supplemental) Summary of advantages of Recentrifuge. This figure summarizes the immediate benefits of applying Recentrifuge to a study involving SMS of different but related samples, including negative controls (see Supplemental Figure 5). Recentrifuge generates four different sets of scored charts for each taxonomic level of interest in addition to the scored plots for the raw samples: samples with the control taxa subtracted, the exclusive taxa per sample and the shared taxa with and without control taxa subtracted. This battery of analysis and plots permits robust comparative analysis of multiple samples in low microbial biomass metagenomic studies.

## 2 Computing kernel implementation details

The following list abridges some implementation details concerning the Recentrifuge computing kernel:

1. Coded using python multi-platform parallelization to reduce the elapsed time when dealing with massive datasets. Depending on the algorithm, the code parallelizes by input file, by taxonomic level or by derived sample —in the summary generation step.
2. Intensive use of recursive methods to cope with tree-arithmetics, which enables robust comparison between taxonomic trees at any rank. The code recognizes the 30 different taxonomic levels used in the NCBI taxonomy (NCBI 1988b). Recentrifuge can adequately deal with arbitrary taxonomic levels (NO_RANK) in between the named NCBI taxonomic ranks, as happens with some complex eukaryotic taxa.
3. Software engineered with robustness as one of the leading targets. It is a full statically annotated Python 3.6 code. PEP-8 (Rossum et al. 2013), PEP-484 (Rosum et al. 2015), and PEP-526 (R González et al. 2016) compliant. Written following the Google python style guide (Patel et al. 2017). Pylint and mypy checked.
4. Implemented under an object-oriented paradigm to ease future extensions targeting new or improved uses, Recentrifuge can be easily extended to understand additional input formats and other taxonomies different from NCBI —by direct support extending the base class or indirectly by using a mapping software like CrossClassify (Balvociüté and Huson 2017). Supplemental Figure 9 Figure shows a summarized UML (Unified Modeling Language) class diagram of Recentrifuge that exposes the main classes developed and currently used in the package.

It is worth to mention that the code allows applying a different parameter set to the control samples, including mintaxa and minscore. This feature is helpful when the control samples are too different from the ordinary ones and thus require unique values for the parameters that define how Recentrifuge treats the sample data.

Supplemental Table 1 shows basic code layout statistics about Recentrifuge (source files, lines, and number of code lines equivalent in third generation languages) provided by cloc (v.1.76).

**Figure 9.**
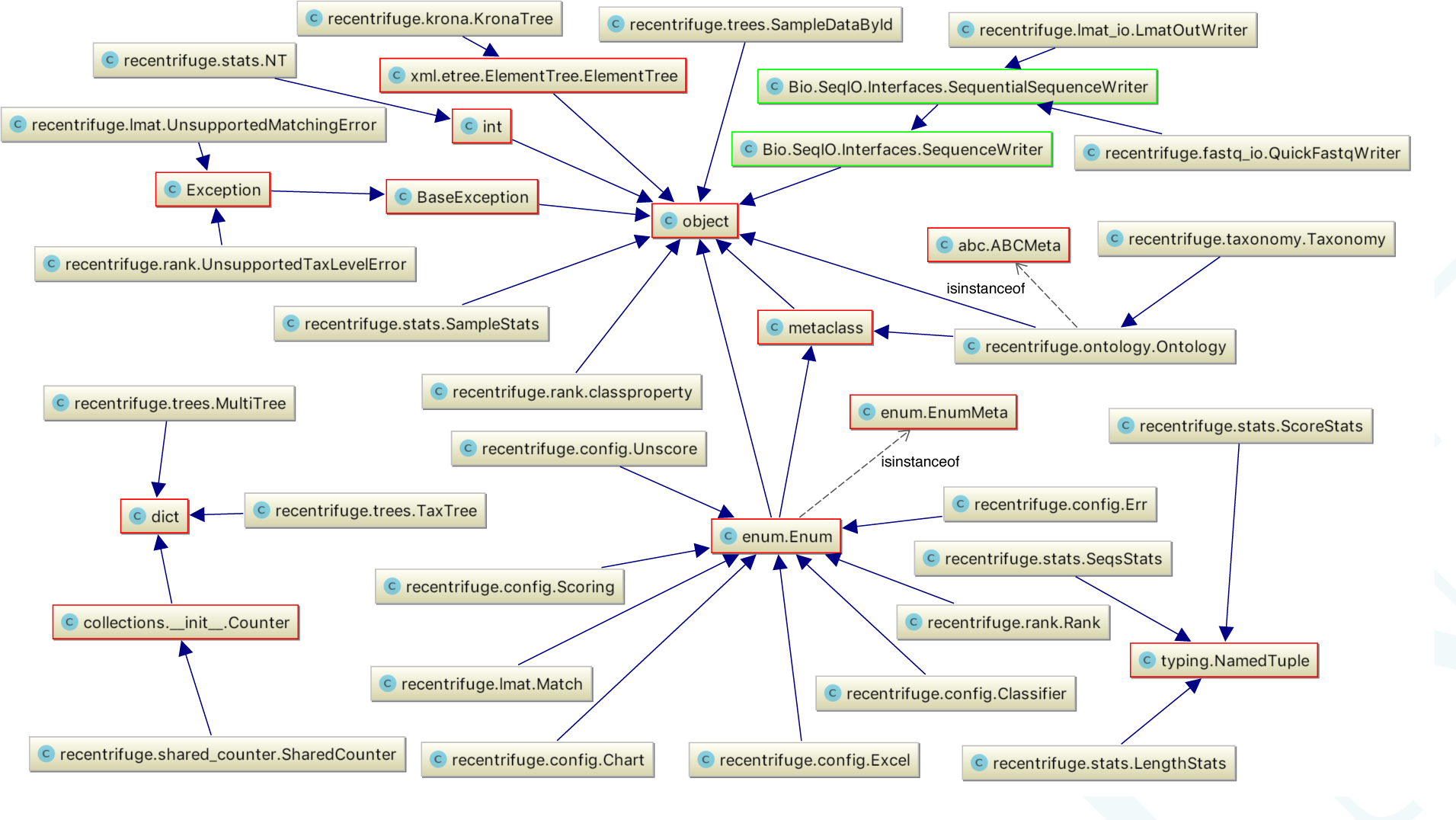
(Supplemental) UML class diagram of Recentrifuge kernel. This Figure summarizes the relationships between developed classes in the Recentrifuge core package. The classes in the figure with a colored border are the parent classes from which the Recentrifuge ones derive. Those belong to the Python Standard Library (red border) and BioPython (green border).

**Table 1.**
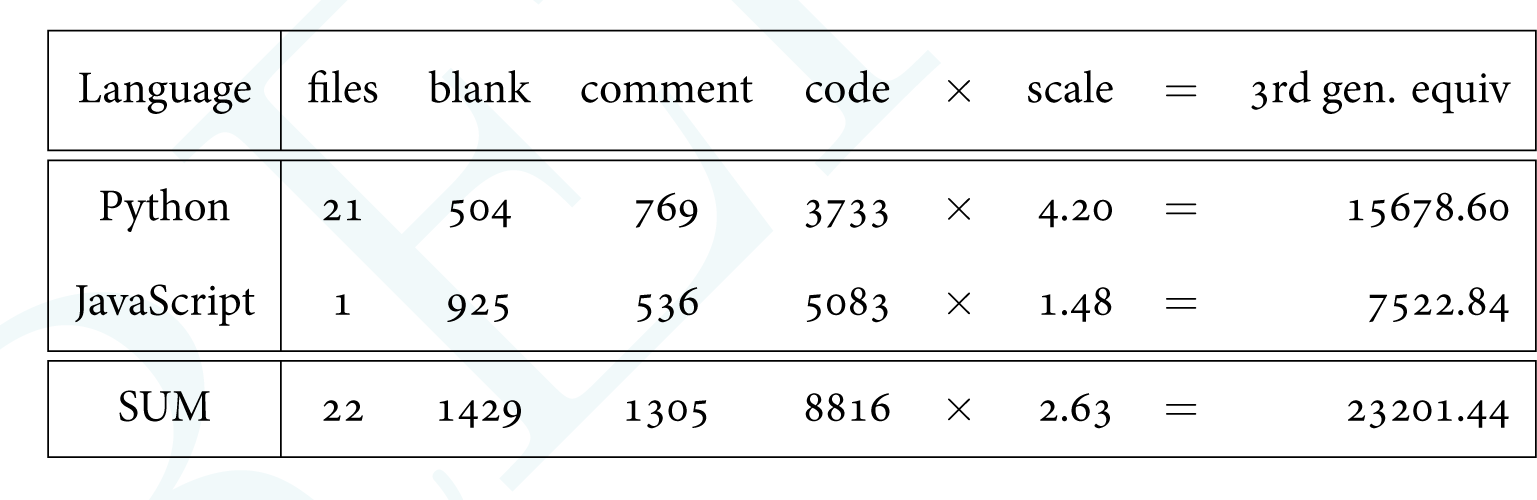
(Supplemental) Recentrifuge code layout. Basic code layout statistics about Recentrifuge (source files, lines, and number of code lines equivalent in third generation languages) provided by cloc (v.1.76)

